## Supplementary figures and images for "Prevalence of *Campylobacter* and non-typhoidal *Salmonella* along broiler chicken production and distribution networks, Vietnam"

### S1 Fig.

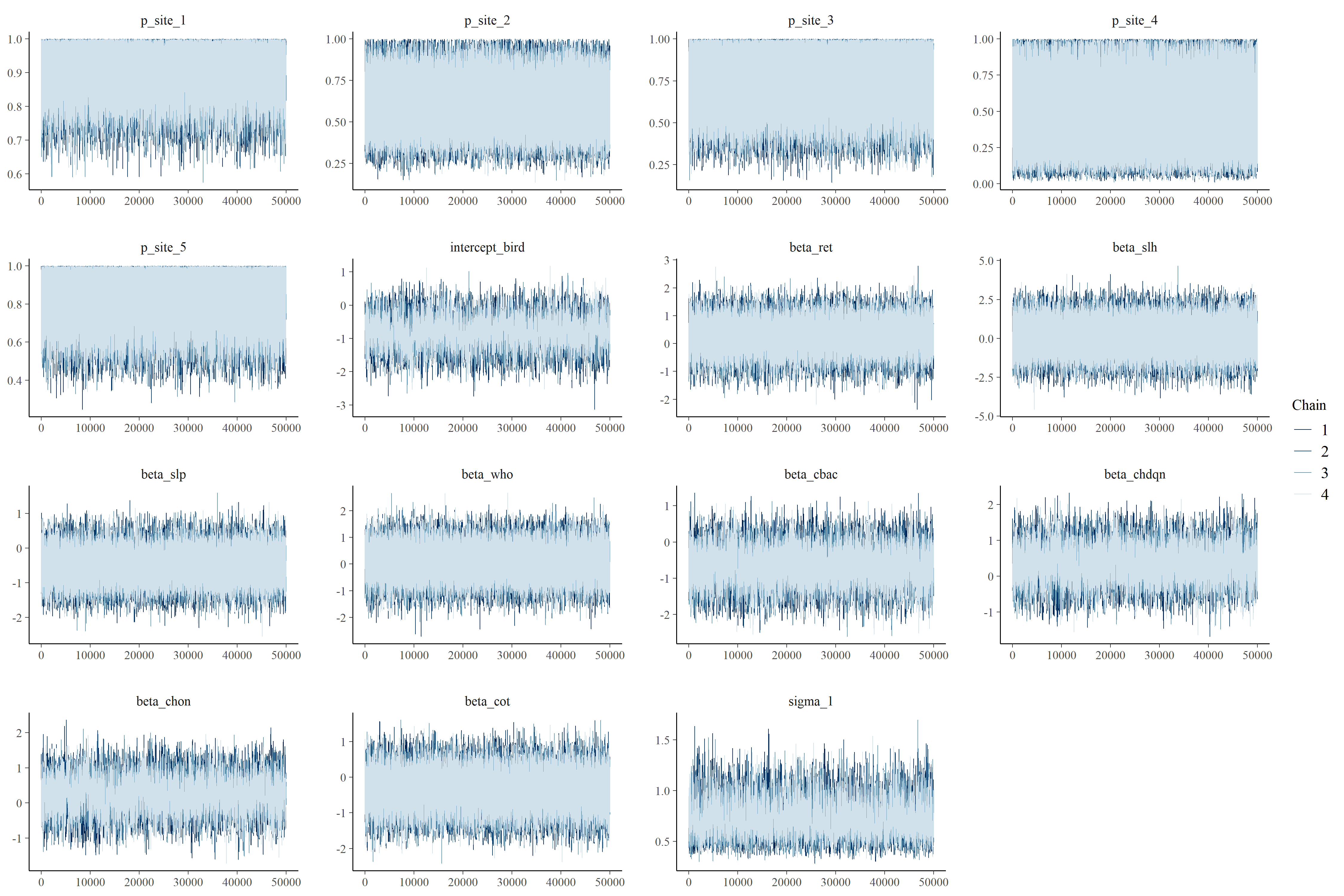

### S2 Fig.

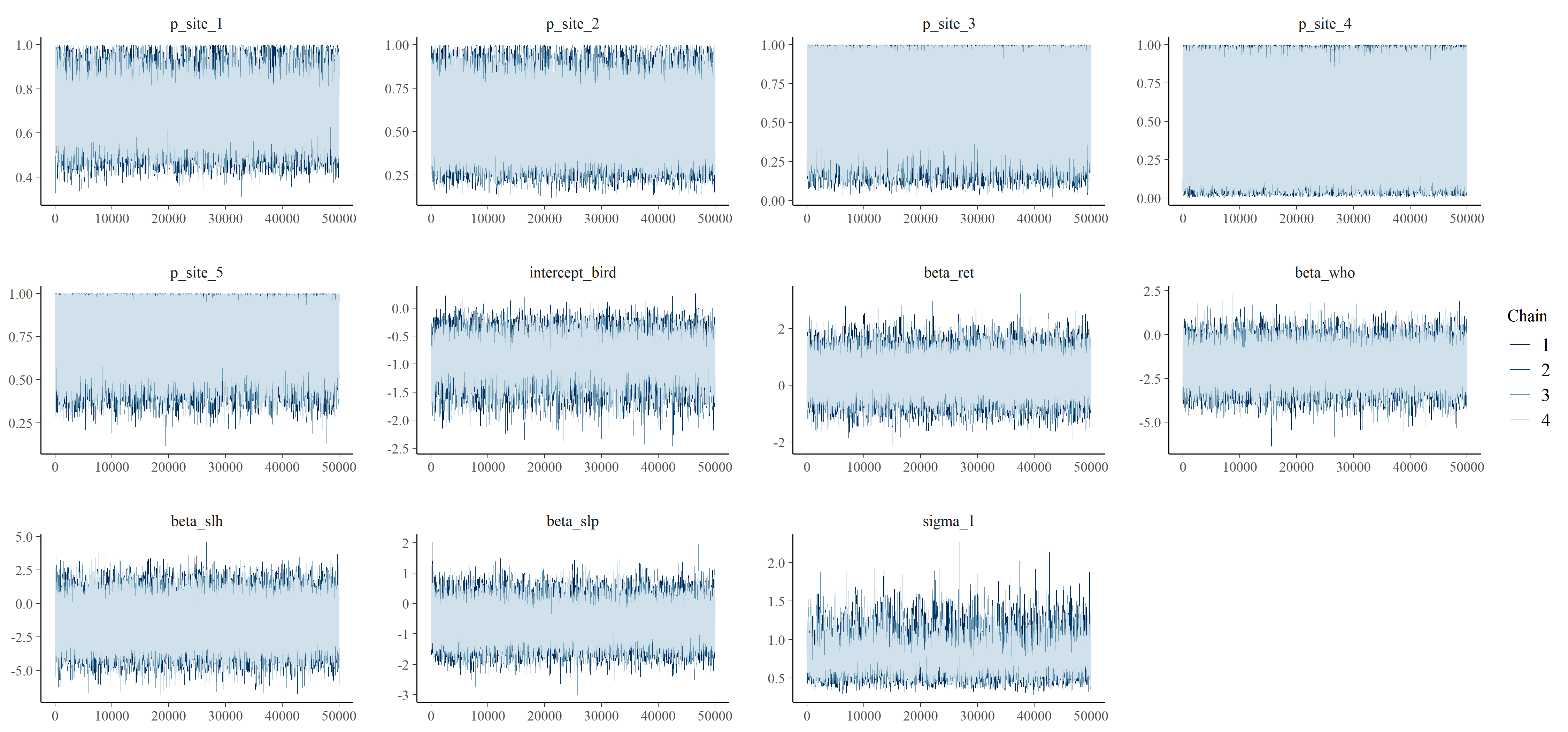

### S3 Fig.

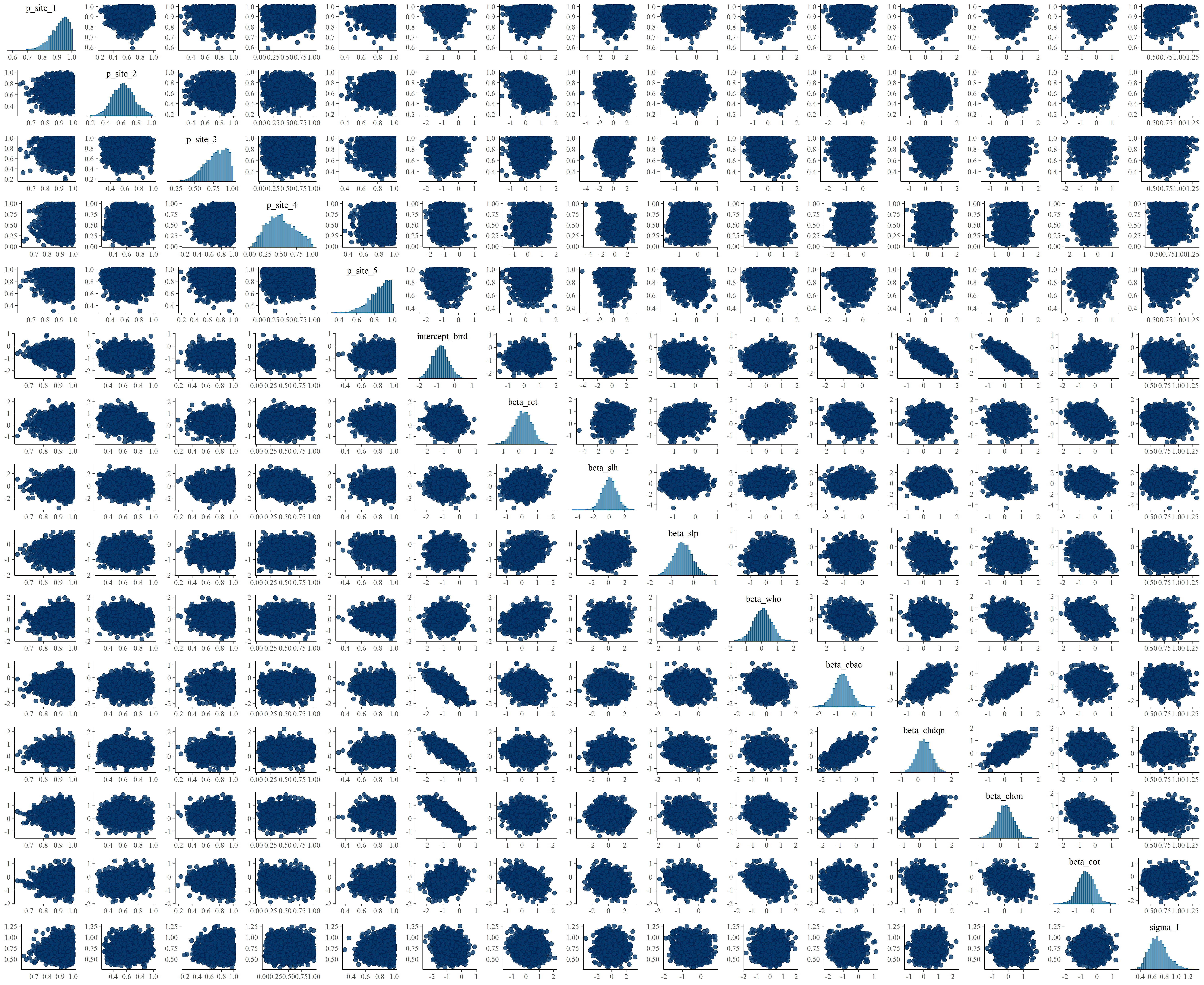

### S4 Fig.

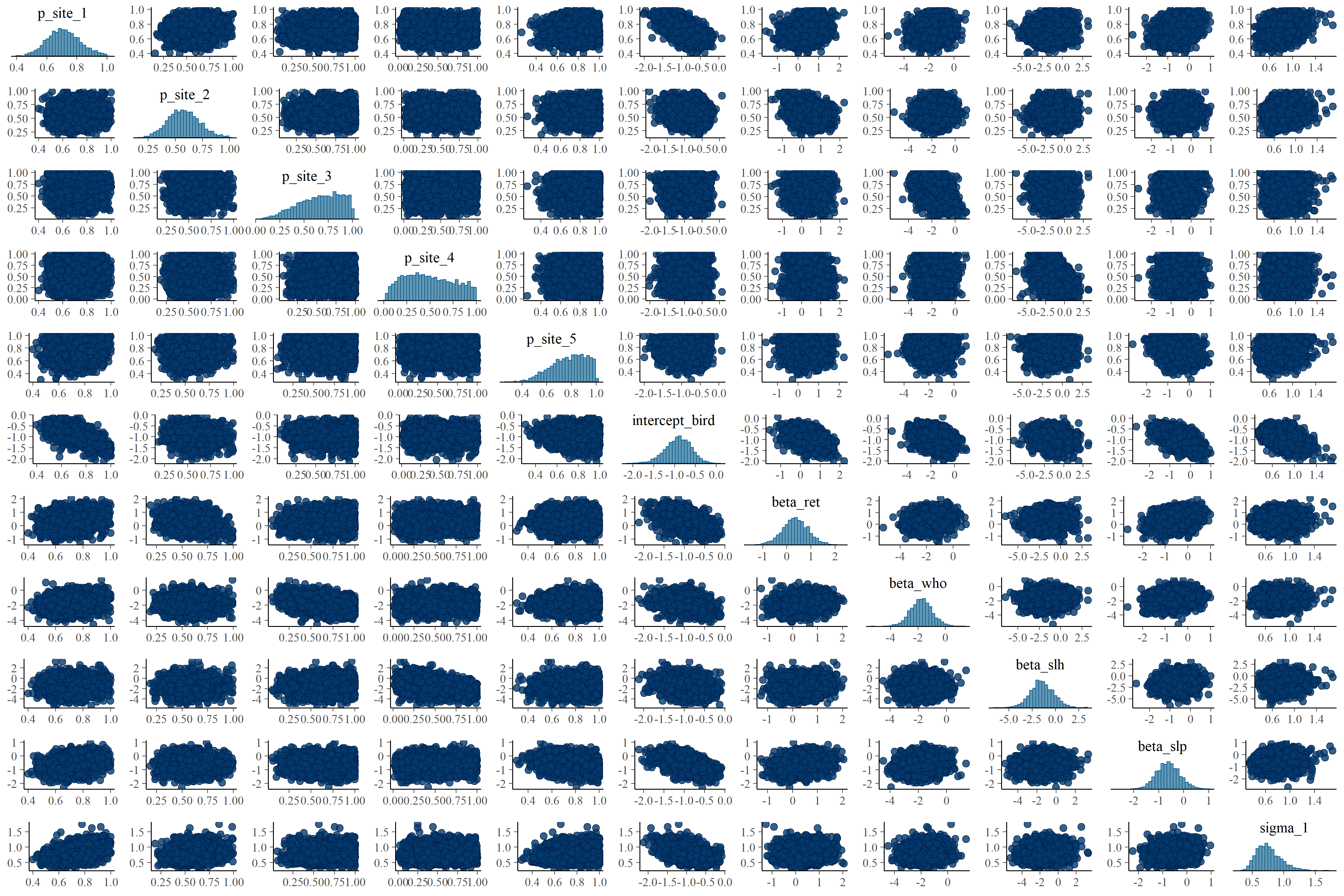

### S5 Fig.

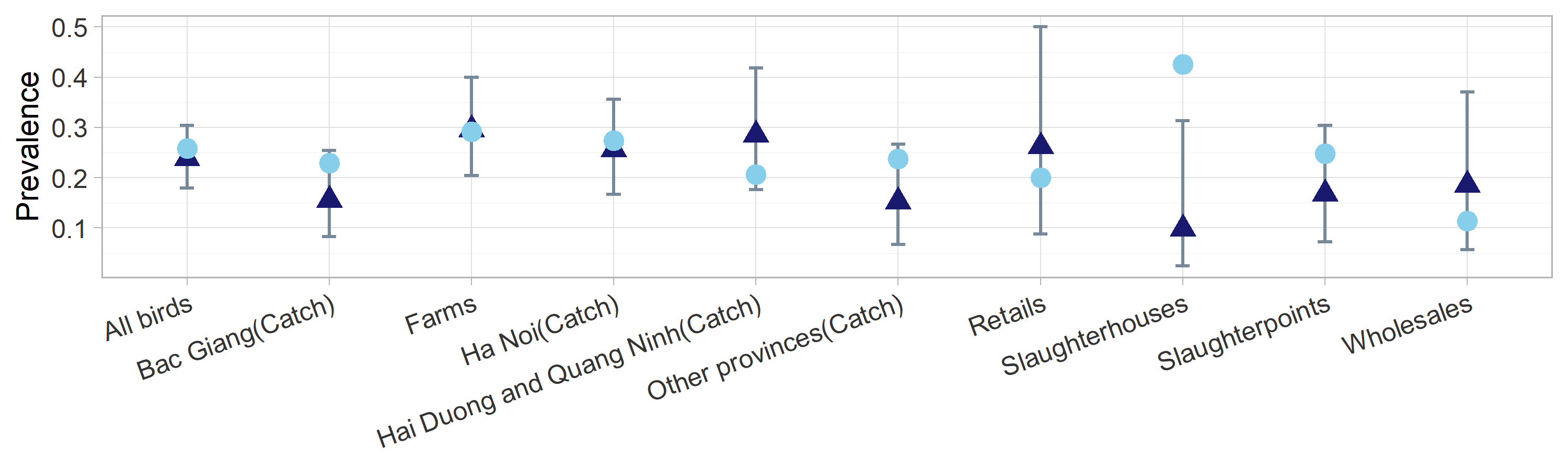

### S6 Fig.

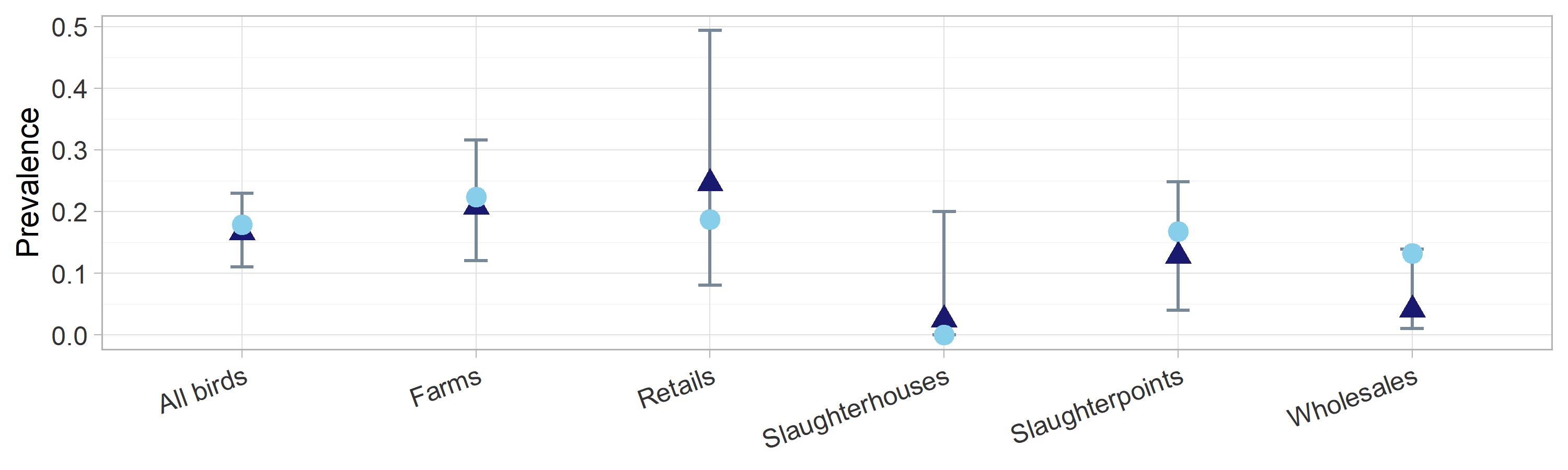

### S7 Fig.

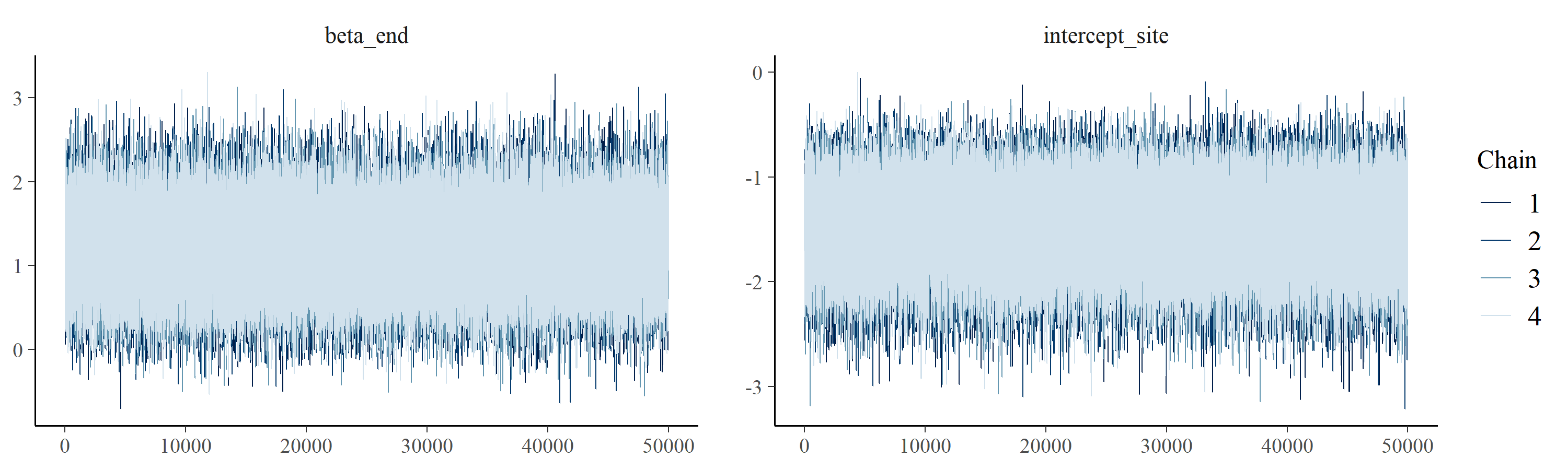

### S8 Fig.

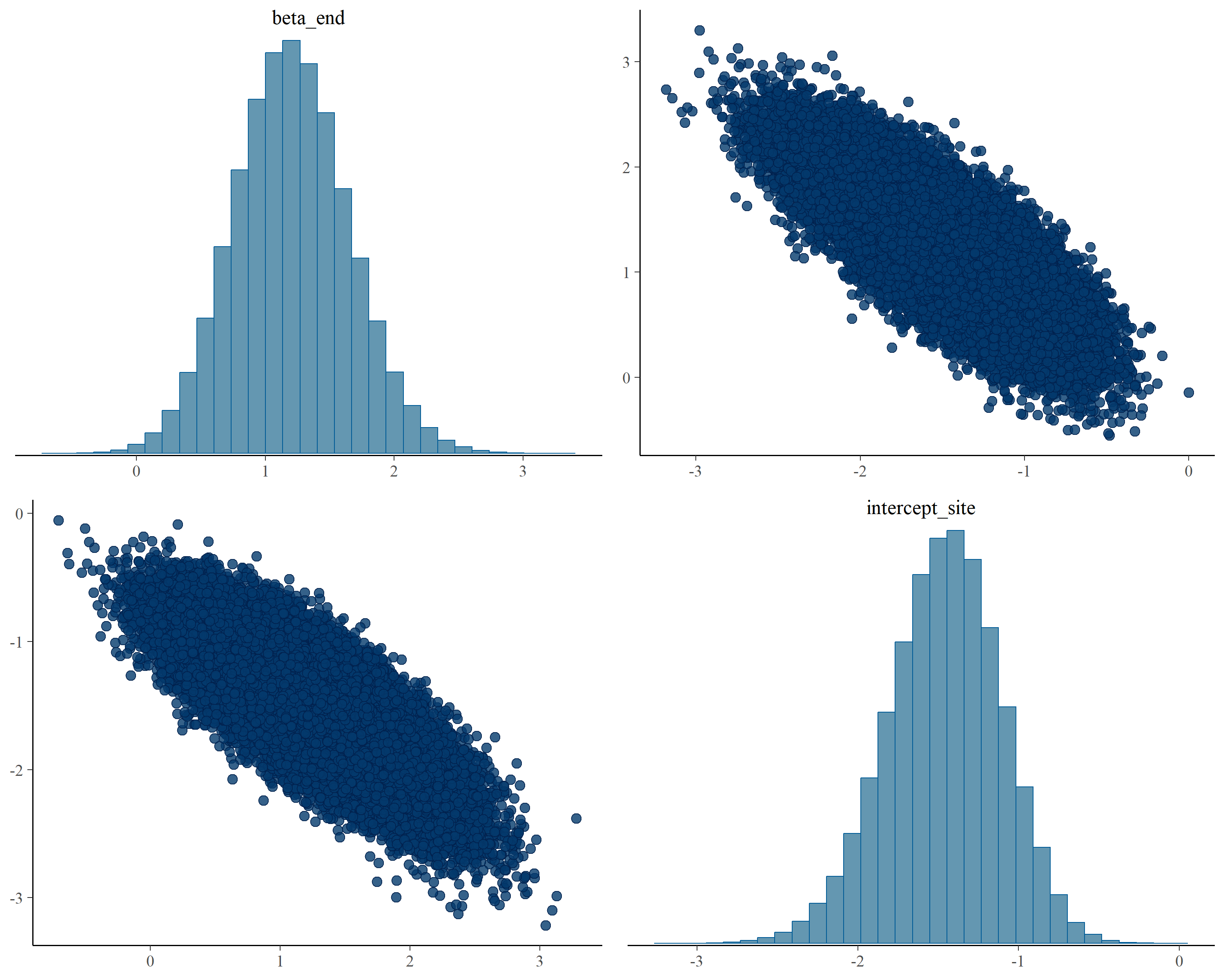

### S9 Fig.

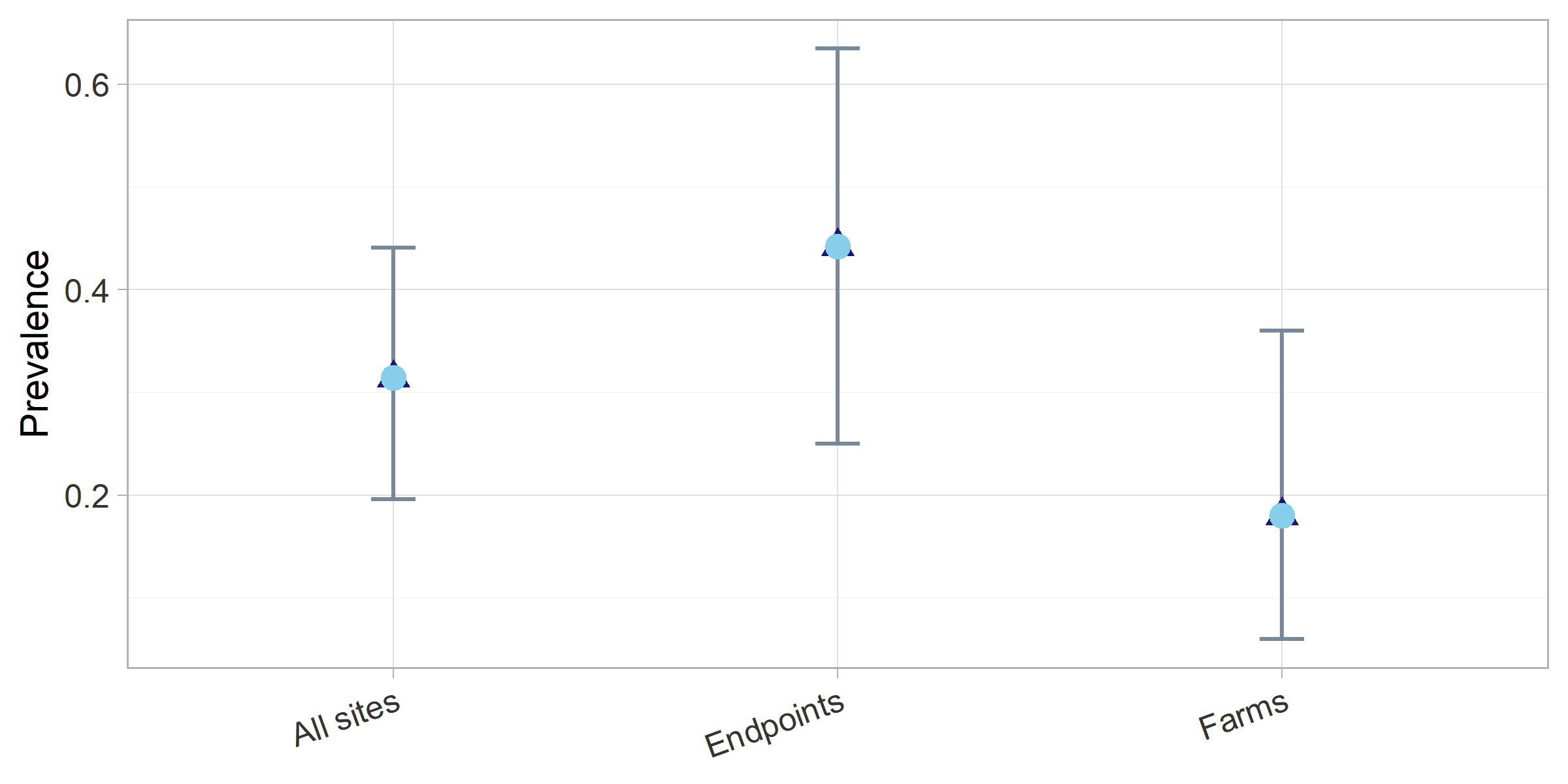
